# Blue light promotes ascorbate synthesis by deactivating the PAS/LOV photoreceptor that inhibits GDP-L-galactose phosphorylase

**DOI:** 10.1101/2022.12.12.520143

**Authors:** Céline Bournonville, Kentaro Mori, Paul Deslous, Guillaume Decros, Tim Blomeier, Jean-Philippe Mauxion, Joana Jorly, Stéphanie Gadin, Cédric Cassan, Mickael Maucourt, Daniel Just, Cécile Brès, Christophe Rothan, Carine Ferrand, Lucie Fernandez-Lochu, Laure Bataille, Kenji Miura, Laure Beven, Matias Zurbriggen, Pierre Pétriacq, Yves Gibon, Pierre Baldet

**Author notes:** Monsanto SAS, 1050 Route de Pardies, 40300 Peyrehorade, France. C. Bournonville and K. Mori contributed equally to this work. The author responsible for distributing materials integral to the findings presented in this article in accordance with the policy described in the Instructions for Authors (www.plantcell.org) is Pierre Baldet. Corresponding author: Pierre Baldet.

## Abstract

Ascorbate (vitamin C) is one of the most essential antioxidants in fresh fruits and vegetables. To get insights into the regulation of ascorbate metabolism in plants, a mutant producing ascorbate-enriched fruits was studied. The causal mutation, identified by a mapping-by-sequencing strategy, corresponded to a knock-out recessive mutation in a new class of photoreceptor named PAS/LOV protein (PLP, Solyc05g07020), which acts as a negative regulator of ascorbate biosynthesis in tomato. This trait was confirmed by CRISPR/Cas9 gene editing, and further found in all plant organs, including fruit that accumulated 2-3 times more ascorbate than in the WT. The functional characterization revealed that PLP interacted with the two isoforms of GDP-L-galactose phosphorylase (GGP), known as the controlling step of the L-galactose pathway of ascorbate synthesis. The interaction with GGP occurred in the cytoplasm and the nucleus, but was abolished when PLP was mutated. These results were confirmed by an optogenetic approach using an animal cell system, which additionally demonstrated that blue light modulated the PLP-GGP interaction. Assays performed *in vitro* with heterologously expressed GGP and PLP showed that PLP is a non-competitive inhibitor of GGP that is inactivated after blue light exposure. This discovery sheds light on the light-dependent regulation of ascorbate metabolism in plants.

## INTRODUCTION

Ascorbate is an essential metabolite in living organisms. It has a leading role as antioxidant, by participating in eliminating reactive oxygen species (ROS) that are usually produced in response to biotic and abiotic stresses (Decros et al., 2019). Ascorbate also plays a crucial role in controlling the levels of ROS that are continuously produced under optimal conditions by cell metabolism, in particular photosynthesis. Due to its high antioxidant potential, ascorbate is one of the most important traits for the nutritional quality of fruits and vegetables. Indeed, evolution in humans and a few animal species has led to the loss of the L-gulono-γ-lactone oxidase activity, which catalyzes the last steps of the biosynthetic pathway (Burns, 1957). Consequently, humans are unable to synthesize ascorbate, thus defined as vitamin C, and must have a daily intake through the consumption of fruits and vegetables. Paradoxically, the domestication of various fruit species has resulted in decreased ascorbate content (Gest et al., 2013), suggesting the occurrence of a trade-off between fruit yield and quality. Thus, understanding ascorbate metabolism is a critical issue in plant breeding, particularly for fleshy fruit species such as tomato, which is considered one of the major sources of vitamin C in the human diet (Wheeler et al., 1998).

The map of plant ascorbate metabolism is well established since the discovery of the Smirnoff-Wheeler pathway, also called L-galactose pathway, although little is known about the regulatory mechanisms involved (Wheeler et al., 1998, Bulley and Laing, 2016). The GGP protein, also known as VTC2 by analogy with *Arabidopsis thaliana*, corresponds to a GDP-L-galactose phosphorylase (Linster et al., 2008), and is so far considered the controlling enzyme of the L-galactose pathway (Bulley, 2009; Li et al., 2013; Fenech et al., 2021). It catalyzes the first step in ascorbate biosynthesis in plants, *i.e.* the conversion of GDP-L-galactose into L-galactose-1-phosphate. Arabidopsis knock-out *vtc2* mutants display a drastic decrease in ascorbate, although a residual amount is still produced (Dowdle et al., 2007), due to the presence of another gene encoding a GDP-L-galactose phosphorylase, namely *VTC5*. The *VTC5* gene has a high sequence homology (∼66% identity) with its counterpart *VTC2,* but it was found 100 to 1000 times less expressed. Other studies have hypothesized the existence of additional alternative pathways (Wheeler et al., 2015). Among them, only the galacturonate and myo-inositol pathways were considered relevant. However, these alternative routes have not been completely demonstrated, and some results tend to invalidate these assumptions. Indeed, the *Arabidopsis vtc2/vtc5* double mutant is unable to grow without the addition of exogenous ascorbate (Dowdle et al., 2007). This suggests that there would be no other way than the L-galactose pathway to complement ascorbate deficiency. Among the enzymes involved in ascorbate synthesis, VTC2 is the only one to have a significant effect on ascorbate levels when overexpressed, although GDP-D-mannose 3’,5’-epimerase acts synergistically with it to increase ascorbate in leaves (Bulley et al., 2009. Precise information on the regulation of VTC2 is lacking, with the exception of the activity of an uORF (upstream Open Reading Frame) in the 5’-UTR of the *VTC2* gene, which was found to control the level of translation of the VTC2 protein (Laing et al., 2015). Interestingly, in the presence of high ascorbate concentration, the VTC2 protein was shown to be downregulated (Laing, 2015). This is to be linked to the fact that excess ascorbate can have deleterious effects, in particular male sterility (Deslous et al., 2021). At the cellular level, a fluorescent fusion protein approach emphasized that the VTC2 protein is localized in both cytoplasmic and nuclear compartments (Müller-Moulé, 2008). This unexpected nuclear localization for a metabolic enzyme suggests that GGP might also act as a dual-function protein: a regulatory factor as well as a catalytic enzyme.

Ascorbate levels are highly dependent on environmental conditions, *e.g.* salt stress, drought and, in particular, intense light that induces ascorbate accumulation. The existence of regulators has been recently demonstrated with the discovery of a few proteins acting at the transcriptional or post-transcriptional level on the regulation of specific genes and enzymes of the L-galactose pathway. These studies, mainly carried out in Arabidopsis leaf, identified AMR1, ERF98 and CNS5B proteins as positive or negative regulators. However, for some of these effectors, the underlying mechanisms remain to be depicted (Zhang and Huang, 2010; Zhang et al., 2012; Alimohammadi et al., 2012). Regarding the effect of light on plant development, previous works showed that light might directly or indirectly affect the expression of genes of the L-galactose pathway. For instance, darkness in Arabidopsis induced the degradation of GDP-mannose pyrophosphorylase (VTC1) by the CSN5B-Cop9 complex associated with the proteasome (Wang et al., 2013). Moreover, the AMR1/SCF complex has been shown to modulate the expression of all genes of the L-galactose pathway through an unknown mechanism (Zhang et al., 2009). Interestingly, prolonged exposure to light increases the expression of the *GGP* gene (Dowdle et al., 2007). Additionally, some studies have shown that the *GGP* gene is under circadian control (Tabata et al., 2002; Dowdle et al., 2007; Müller-Moulé, 2008). All these studies demonstrate an apparent link between ascorbate metabolism and light signalling, but to date there is no evidence that light directly induces ascorbate biosynthesis or that overexpression or activation of the GGP enzyme is the consequence of oxidative stress related to light exposure.

It is well-established that there is a positive correlation between light intensity and ascorbate levels in photosynthetic tissues (Gautier et al., 2008; and Bartoli et al., 2009). In tomato, light was also found to impact ascorbate content more in the leaves than in fruit (Massot et al., 2012). Furthermore, Gautier et al., (2009) showed that fruit ascorbate content was not limited by leaf photosynthesis but was dependent on direct fruit irradiance. Surprisingly, the literature is poor regarding the molecular mechanisms relating ascorbate and light sensing. The photoreceptor proteins that collect a light signal *via* the absorption of a photon to drive and govern biological activity are classified according to the wavelength and physiological processes involved. Among these, phytochromes (red and far-red) and UVR8 (UVB) play a crucial role in plant development and UV protection. The largest family includes the blue light photoreceptors, which are involved in the circadian clock and phototropism through protein and gene expression modifications (Christie, 2007). These blue light sensors are all flavoproteins, thus requiring a flavin (FMN or FAD) cofactor domain in addition to the effector domain. Most of them are well characterized, such as the phototropins and the proteins involved in the circadian clock (Briggs, 2001; Christie et al., 2002; Crosson et al., 2003). Recent reviews mentioned another type of photoreceptor protein containing a unique LOV (Light Oxygen Voltage) domain, named PAS/LOV Protein (PLP), and for which the biological function remains to be established. It has been demonstrated that the LOV domain of these photoreceptor proteins can change conformation when exposed to blue light and that this modification is reversible in the absence of blue light signal (Kasahara 2010). Recently, Li and co-workers (2018) showed that the expression of PLP can also be triggered in soybean plants cultured under darkness or red light. Interestingly, a yeast two-hybrid screening performed with a cDNA bank from Arabidopsis leaf allowed the identification of AtGGP1 (VTC2) and AtGGP2 (VTC5) as being potential PLP-interacting proteins (Ogura et al., 2008). In their study carried out in soybean, Li and co-workers observed some phenotypic alterations in the *Gmplp* mutants, especially hypocotyl growth. Nevertheless, no hypothesis was proposed to explain the putative function of the PLP protein in relation to such developmental processes (Li et al., 2018).

A population of EMS mutants obtained in the miniature Micro-Tom tomato cultivar was recently shown to exhibit a genetic and phenotypic variability far beyond the natural variation found in domesticated species (Just et al., 2013; Garcia et al., 2016). In a forward genetic strategy, this population proved valuable to screen for traits of interest (Garcia et al., 2016), including increased ascorbate (Deslous et al., 2021). The present study found a mutation causing a build-up in fruit ascorbate content and validated it within the gene encoding PLP. The interaction of PLP and GGP was confirmed *in vivo* and established *in vitro*, revealing that PLP inhibited GGP and that light promoted ascorbate synthesis by counteracting this inhibition.

## RESULTS

### Detection of ascorbate-enriched mutants in an EMS Micro-Tom Tomato population

A forward genetic approach was performed to discover new regulators of ascorbate metabolism in plants. In that aim, the EMS (Ethyl Methanesulfonate) mutant collection generated in the Micro-Tom Tomato cultivar at INRAE Bordeaux (Just et al., 2013) was screened to find mutants producing ascorbate-enriched (also called AsA+) fruits (Deslous et al., 2021). A total of 500 M2 and M3 mutant families were cultivated in the greenhouse, in batches of 100 families representing a total of 6,000 plants screened. On each plant, at least 4 fruits at the red ripe stage were pooled and assayed for ascorbate content. The range of ascorbate content found in the mutants varied from 0.5 to 4 µmolµmol.g^-1^ FW (Suppl. Fig.1A), whereas the WT mean values ranged from 1 to 1.3 µmolµmol.g^-1^ FW depending on the period of the year. Although low, such variations in ascorbate content were expected in the WT, as it is well known that ascorbate content is regulated by environmental factors such as light irradiance (Gatzek et al., 2002; Gautier et al., 2008; Bartoli et al., 2009).

An ascorbate threshold value for selecting ascorbate-enriched mutants was set at 2 µmolµmol.g^-1^ FW *i.e.* around twice the value of the WT, resulting in the selection of 93 families (193 plants in total). Selected plants were cut to allow new growth, then used in a second screening for confirmation of the “AsA+” phenotype. Following this second screening, 5 families with an ascorbate fruit content 3 to 5 times higher than that of WT were selected. (Suppl. Fig 1B). We present here the characterization of one of these mutants, named P21H6. For this family, only one plant over 12 sown displayed the ascorbate-enriched phenotype. Interestingly, ascorbate content was 2.5 µmolµmol.g^-1^ FW at the first screening performed at the end of autumn, whereas after the confirmation performed during the following spring season, the ascorbate content increased to up to 4 µmol.g^-1^ FW, thus suggesting an impact of the season on ascorbate pools. The P21H6 mutant was chosen as there was no apparent alteration in the phenotype of the vegetative and reproductive organs (Suppl. Fig.1B).

### Identification of the causal mutation of the ascorbate-enriched P21H6 mutant

In order to identify the mutated locus responsible for the ascorbate-enriched phenotype, the genetic inheritance features of the mutation were first determined using classical Mendelian genetics. For this, the ascorbate-enriched phenotype was analyzed in the progeny after self-pollination (S1). For 12 S1 plants, all produced ascorbate-enriched fruits, whereas fruits from all plants of the backcross (BC_1_F_1_) displayed a WT-like ascorbate content phenotype (Suppl. Fig.2A). The resulting BC_1_F_2_ segregating population consisting of 441 individuals was then analyzed for the ascorbate-enriched phenotype (Suppl. Fig. 2B). The analysis showed that 115 plants were defined as “AsA+” and 326 as ‘WT-like’, thus confirming a Mendelian 1:2:1 segregation, involving a single recessive mutation.

For the identification of the causal mutation, a mapping-by-sequencing strategy was used (Garcia et al., 2016). From the BC_1_F_2_ population, two bulked pools of 44 individual plants displaying either AsA+ or WT-like fruit phenotypes were generated (Suppl. Fig.2B). Pooled genomic DNA from each bulk was then sequenced to a tomato genome coverage depth of 39X, the trimmed sequences were mapped onto the tomato reference genome, and EMS mutation variants were filtered to exclude natural polymorphisms found in cv Micro-Tom (Kobayashi et al., 2014) compared with the cv Heinz 1706 reference genome (Suppl. Table 1 and 2). Analysis of the allelic frequencies (AF) of variants in the two bulks led to the identification of chromosome 5 as the genome region carrying the causal mutation since it displayed high mutant AFs (AF > 0.95) in the AsA+ bulk and much lower frequencies (AF < 0.4) in the wild-type-like bulk (Fig.1A). Analysis of the putative effects of the mutations on protein functionality highlighted 2 genes carrying mutations in exons. Among these, one affected an “unknown protein”, while the second affected a predicted PLP (Solyc05g007020) that was located at the top of the SNPs plot according to AF analysis. Given that a link between PLP and the enzyme GGP had previously been found (Ogura et al., 2008), we considered PLP to be the most likely candidate. Unequivocally, to associate the *plp* mutation with the ascorbate-enriched trait, and to exclude any other mutated locus, recombinant plants selected from the BC_1_F_2_ progeny were used (Fig.1B). To this end, we used the EMS-induced SNPs surrounding the *PLP* gene as genetic markers. It clearly indicated that a single G to A nucleotide transversion occurred in the fifth exon corresponding to the LOV domain of PLP (Fig.1C), resulting in the knocking-out of the protein due to the appearance of a STOP codon instead of a glutamine codon.

**Figure 1.**
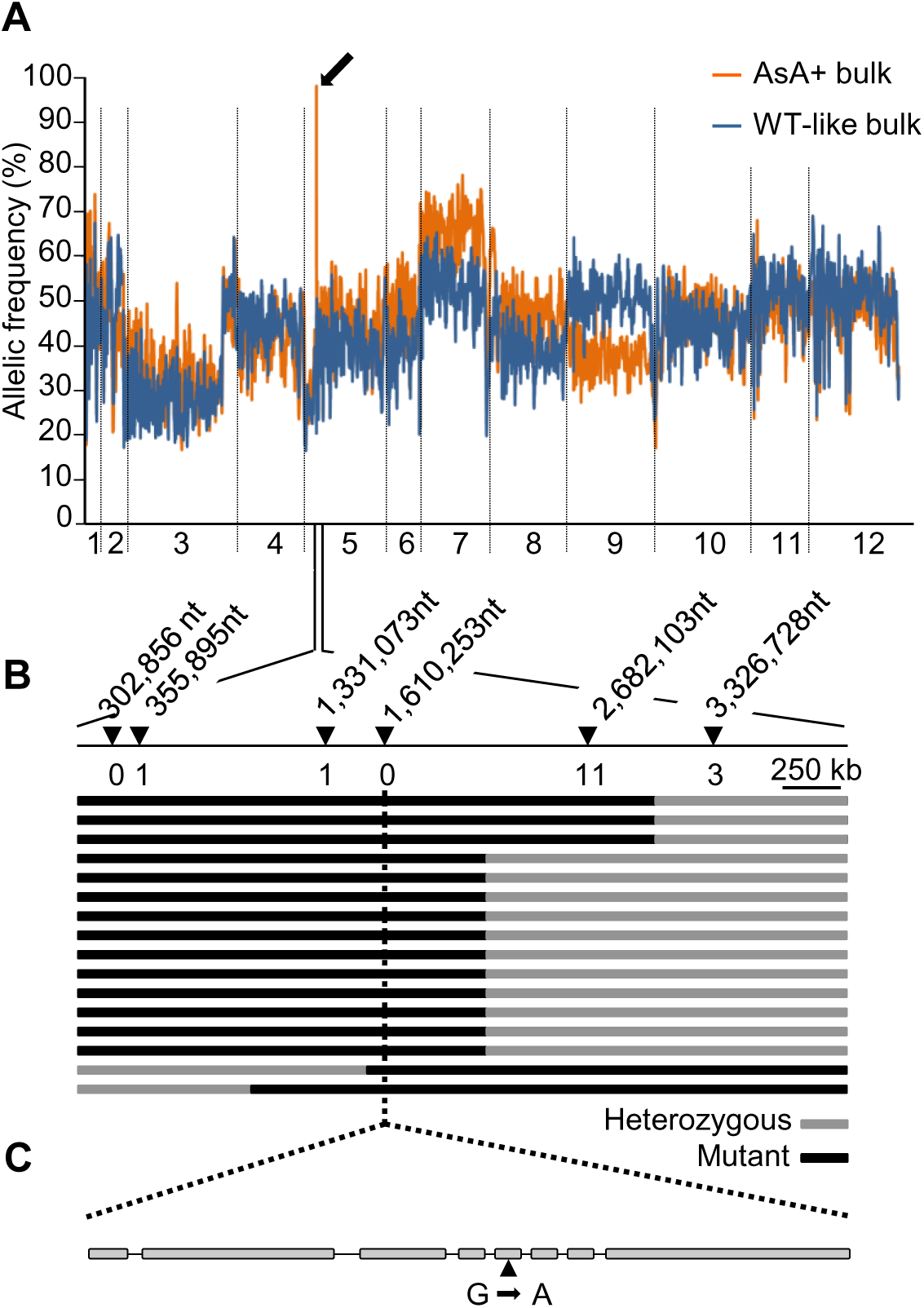
Identification of the mutation responsible for the ascorbate-enriched fruit phenotype. A. Identification of the chromosome associated with the ascorbate-enriched phenotype. Pattern of the mutation allelic frequencies obtained in the mutant and WT-like bulks are represented along tomato chromosomes by black and grey lines, respectively. The plot represents allelic frequencies (*y* axis) against genome positions (*x-*axis). A sliding window of 5 SNPs was used. The *x*-axis displays the 12 tomato chromosomes, the black arrow indicates the peak of allelic frequency (AF) of the chromosome 5 region carrying the putative causal mutations, since it displayed an AF > 0.95 (orange line) in the AsA+ bulk and an AF < 0.4 (grey line) in the WT-like bulk. B. Fine mapping of the causal mutation using the BC_1_F_2_ population. Recombinant analysis of 44 BC_1_F_2_ individuals displaying the ascorbate-enriched phenotype allowed us to locate the causal mutation at position 1,610,253 nucleotides. Black triangles indicate marker positions. Number of recombinants are shown below the position of the markers. Chromosomal constitution of the recombinants is represented by black and grey bars, for mutant and heterozygous segment respectively. C. A single nucleotide transversion, G to A at position 1,610,253 in the Solyc05g007020 fifth exon sequence led to a STOP codon.

The functional validation of PLP as a negative regulator of ascorbate biosynthesis was next performed by generating *plp* mutants in the WT using Clustered regularly interspaced short palindromic repeat (CRISPR)/CRISPR-associated protein 9 (Cas9). Within the large family of photoreceptor proteins harboring a LOV domain, the latter has a very high level of sequence homology. Consequently, the PAS domain was targeted to produce KO mutations in the 5’ terminal region of the *PLP* gene (Fig.2A). Several T0 lines were generated, and mature fruits were analyzed for ascorbate content. Five T0 lines displaying at least a 2-fold increase in ascorbate were selected to produce the next T1 generation (Suppl. Table 3). The sequencing of the *PLP* gene for several T1 plants of lines 6, 15, 17 and 21 revealed various deletions that represented 1 to 8 nucleotides (Suppl. Fig.3). All of these deletions resulted in a shift of the open reading frame and the appearance of a STOP codon in the downstream coding sequence of the *PLP* gene. For further studies, homozygous T2 plants of lines 15-1 and 15-5 were used indifferently as they harbored the same mutation (Fig. 2B; Suppl. Fig. 3). Moreover, a tomato genome investigation revealed the presence of a PLP-like protein (Solyc01g010480) displaying 65% peptide identity mainly distributed in the PAS and LOV domains characteristic of the phototropin proteins. Sequencing analysis showed that this PLP-like protein was not off-targeted by the chosen CRISPR/Cas9 strategy.

**Figure 2.**
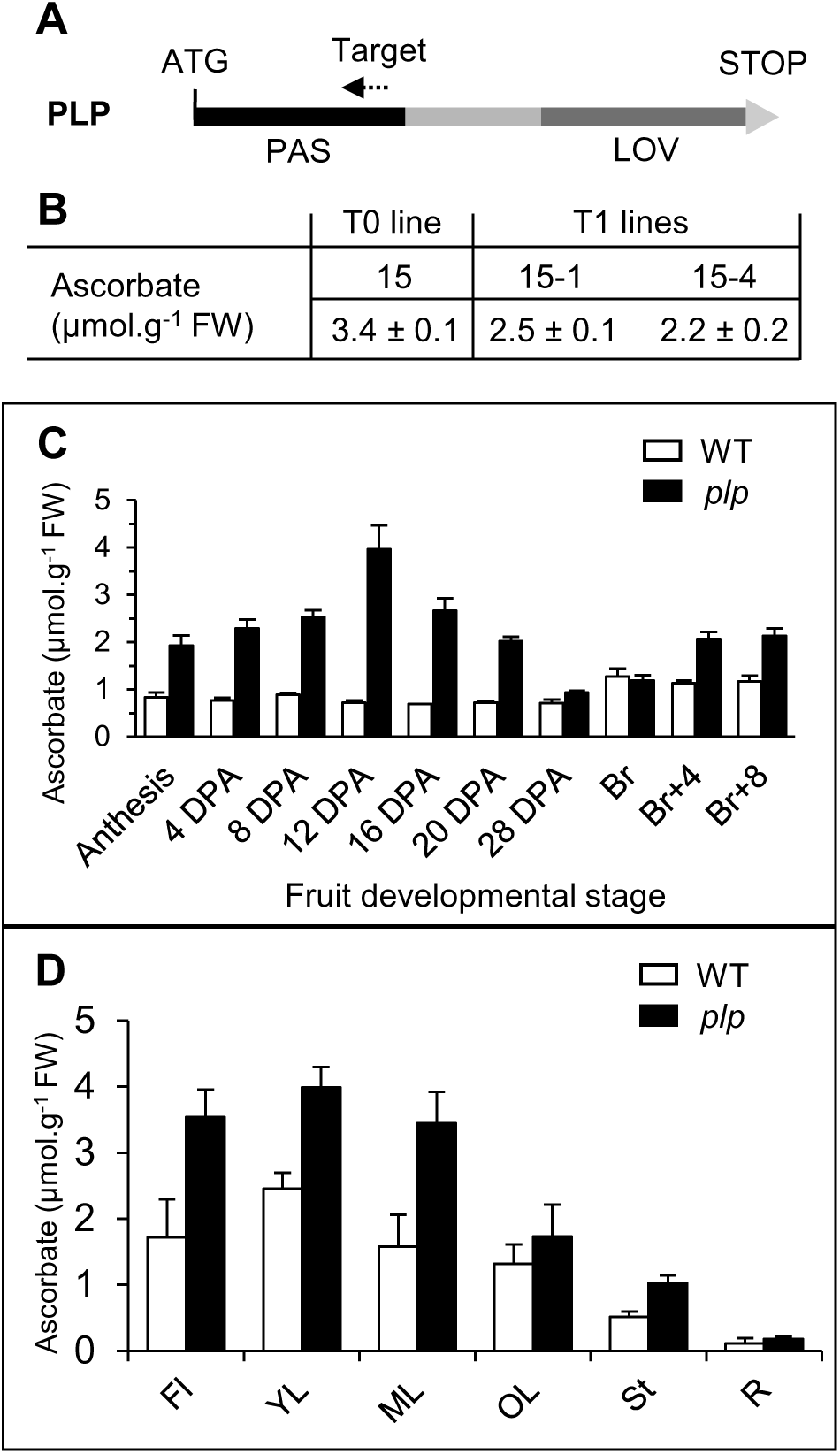
Validation of PAS/LOV as the candidate gene involved in regulating ascorbate content in developing fruit and several tomato plant organs. A. Schematic representation of *PLP* showing its PAS and LOV domains. The dashed arrow in the PAS domain indicates the position of the target sequence for the CRISPR-Cas9 construct. B. Ascorbate in red ripe fruit (means ± SD, n=4) of WT, T0 line 15 and progeny T1 lines 15-1 and 15-4. C. Ascorbate content in fruit of WT and *plp* mutant plants during development, from anthesis to ripeness. D. Ascorbate content in flowers, leaves at 3 stage of development, stem and roots of the 15-5 line and control one-month-old plants. Data are the means ± SD of a total of three individual plants per organ and three organs per plant, except for anthesis (100 organs) and at 4 DPA (20 organs). Abbreviations: PLP, PAS/LOV; DPA, days post-anthesis; Br, breaker stage; FI, flowers; YL, young leaf; ML, mature leaf; OL, old leaf; St, stem; R, roots.

### PAS/LOV is a repressor of ascorbate accumulation in tomato plants

As mentioned above, no noticeable morphological and physiological change was observed at the whole plant level in the mutant compared to the WT. To characterize the consequences of the knocking-out of the *PLP* gene on ascorbate metabolism, ascorbate assays were carried out during fruit development as well as in all tomato plant organs. As shown in Fig. 2C, growing fruits of the *plp* mutant had more ascorbate than the WT, with a peak at 12 DPA. Interestingly, the difference in ascorbate content between mutant and WT declined to become negligible at the beginning of the maturation phase (Breaker stage) but rose again during maturation. Finally, there was more ascorbate in all vegetative and reproductive organs of the mutant (Fig. 2D).

Next, we focused on leaves, which are easier to investigate than fruits. Since the mutated protein is a photosensor, the evolution of ascorbate across a day and night cycle was compared with those of the transcripts encoding PLP and GGP1. Every two hours, photosynthetically active radiation (PAR) and temperature were recorded (Fig. 3A), and plant material harvested. As illustrated in Fig. 3B, while the level of ascorbate was always higher in the mutant, the daily variation in ascorbate content followed the same pattern in the T2 line 15-5 mutant and the WT. Thus, in both genotypes, ascorbate increased or plateaued during the first part of the day, then decreased towards the beginning of the night to increase again during the night. This was corroborated by the changes in *GGP1* mRNA abundance, which decreased during the day and increased during the night, peaking 2 h before sunrise (Fig. 3D). Interestingly, *PLP* mRNA also peaked at 4 h in the night. However, *PLP* transcripts were less decreased during the day than those encoding GGP1. Also, as expected, levels of *PLP* transcripts were significantly lower in the *plp* mutant compared to WT (Fig. 3C).

**Figure 3.**
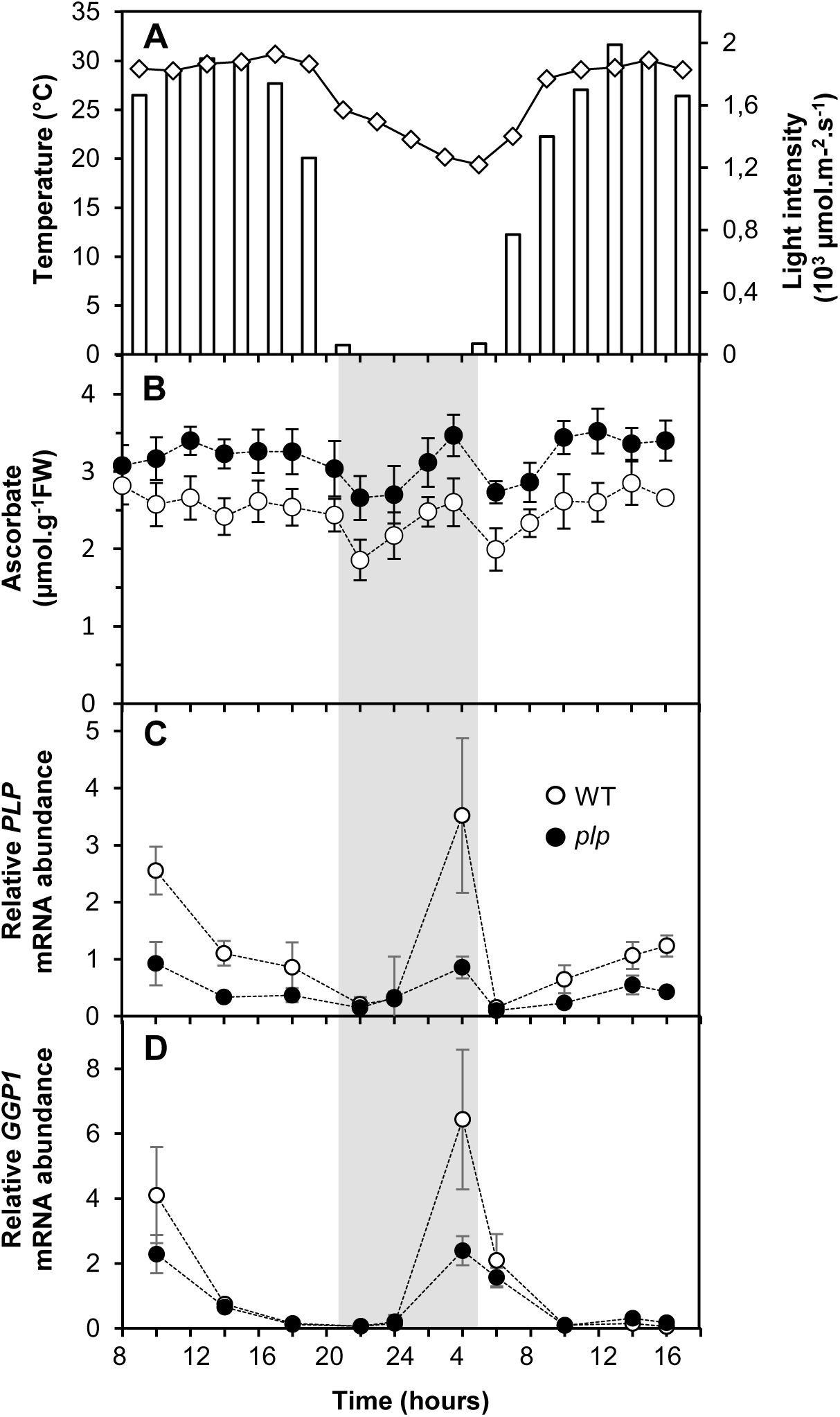
Changes in ascorbate and GDP-L-galactose phosphorylase and PAS/LOV mRNA during a day and night cycle in *plp* mutant and WT plants. *plp* mutant (T2 line 15-5) and WT plants were cultured in the greenhouse for 1 month. The night before the beginning of the experiment, all plants were moved outside and maintained under natural light conditions during 32 hours. This experiment was carried out twice, on May 19^th^ and July 10^th^ 2018, they both lead remarkably to the same results. Here is presented the data obtained on July 10^th^. A. Ambient temperature (diamonds) and light intensity (bars). B. Ascorbate content. C. *PLP* mRNA abundance. D. *GGP1* mRNA abundance. Data are expressed as means ± SD of a total three mature leaves from three individual plants from the 15-5 *plp* line and WT control.

### PAS/LOV interacts with GDP-L-galactose phosphorylase in the cytosol and the nucleus under the dependence of light signaling

In Arabidopsis, previous works have suggested a potential interaction between PLP and VTC2, which was governed by the light spectrum (Ogura et al., 2008). We first studied the subcellular localization of PLP by transient expression in *Nicotiana benthamiana* leaves. For this, a 35S-GFP-PLP construct was used, either under its WT or truncated forms as well as the two GGP isoforms, *i.e.* 35S-GFP-GGP1 and 35S-GFP-GGP2. Confocal microscopy analyses showed that the WT PLP, truncated PLP, GGP1 and GGP2, all localized in the nucleus and the cytoplasm (Fig.4A). The dual localization of GGP1 is consistent with previous findings in Arabidopsis for VTC2 (Müller-Moulé, 2008), which is a homolog to GGP1. However, according to previous literature, the subcellular localization of PLP, and more generally of blue light receptors, still needs to be established. To rule out the leakage of a cleaved GFP from PLP, we performed an LC-MS/MS peptide analysis after SDS-PAGE separation of a crude protein extracts from the stable transgenic tomato harboring the GFP-PLP construct. Peptides derived from the GFP-PLP fusion protein (Suppl. Fig. 4) were detected in the gel band, corresponding to an apparent molecular mass of 73 kDa. This is consistent with the recovery of the non-truncated fused GFP-PLP protein after gel separation. No peptide derived from the GFP-PLP was detected in lower gel bands, in which cleaved products would have been expected to be recovered, confirming the integrity of the fused GFP-PLP after gel separation and, thus, during confocal microscopy analyses. Next, we tested the physical interactions of PLP with GGP1 by using the BiFC technique in simple onion cell, with a combination of vectors for the *GGP1, GGP2, PLP* and mutated *PLP* (hereafter referred to as *PLP^m^*) sequences (Suppl. Table 4). The interaction between the two proteins was confirmed, but not with the truncated PLP^m^ protein (Suppl. Fig. 5). Interestingly, the interaction occurred in both the cytoplasm and the nucleus. Since the mutation is characterized by a truncation of the blue-light sensing LOV-domain, a further verification of the protein-protein-interactions and their light-dependency needed to be tested. For this, a heterologous system was used, thus allowing reconstructing and evaluating of the minimal protein interaction complex upon introduction of the individual components. Here, we implemented mammalian cells, already known for being suitable for expressing plant proteins (Beyer et al., 2015; Müller et al., 2014), and without any additional plant-components that might preclude a straightforward analysis of the interaction. In brief, a tetracycline-based split transcription factor approach (Müller et al., 2014) was customized and used for testing the light-regulated interaction of all GGP and PLP-combinations (*GGP1*, *GGP2*, *PLP* and *PLP^m^* sequences, see Suppl. Table 5). The PLP and PLP^m^ were C-terminally fused to the tetracycline repressor (TetR) that binds to the tetracycline operator (tetO)-motif on the reporter plasmid. GGP1 and GGP2 were either C- or N-terminally coupled to the transactivation domain from the herpes simplex virus type 1 virion protein16 (VP16). Only the interaction of both proteins reconstitutes a functional transcription factor, capable of binding to the tetO-motif in close proximity to the PCMVminimal promoter and inducing gene expression of the reporter gene, *via* the VP16 transactivation domain. A strong interaction was found between WT PLP and both GGP1 and GGP2 in darkness, while the exposure to blue light (455 nm) minimized this interaction in all tested combinations (Fig. 4B). Then, no interaction was found between the PLP^m^ and GGP1 or GGP2, thus confirming the effect of the truncation (Fig. 4B). Interestingly, no additional factor was needed for the interaction of the WT PLP and GGP1 and GGP2. In summary, this experiment clearly showed the light-controlled interaction between PLP and GGP proteins, while the mutant protein completely lost its ability to bind GGP1 or GGP2.

**Figure 4.**
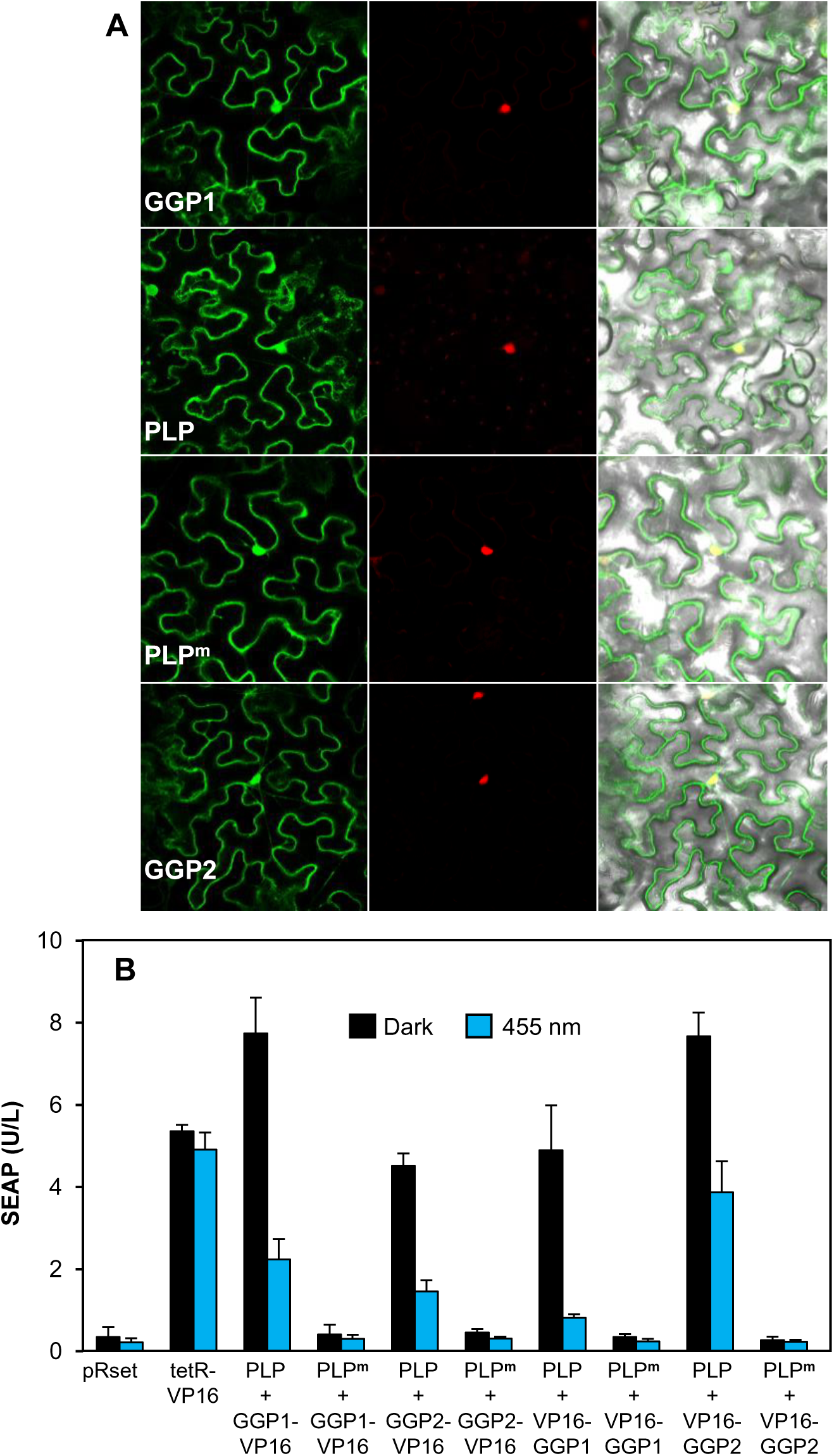
Subcellular localization of PAS/LOV and GDP-L-galactose phosphorylase and their interaction. A. Localization of 35S-GFP-fused proteins in *Nicotiana benthamiana* leaves, co-transformed with nuclear NLS-mcherry as nuclear marker. B. Analysis of PLP-GGP1/2 interactions and its light dependency in a heterologous mammalian split transcription factor system. 50 000 HEK-293T cells were seeded in 24-wells plates and transfected after 24 h with the plasmids pMZ1214, pMZ1215, pMZ1216, pMZ1217, pMZ1218, pMZ1219, pMZ1240, pMZ1241, pSAM, pRSET and pKM006. Twenty-four hours post transfection, the medium was exchanged by fresh medium and the cells were illuminated at 455 nm light (10 μmol m^-2^s^-1^) or kept in the dark for 24 h prior to SEAP quantification Data are represented as means ± SD (n=4). Abbreviations: HEK-293T, human embryonic kidney cells; SEAP, secreted alkaline phosphatase.

### Repression of ascorbate accumulation *via* PAS/LOV is modulated by light

To investigate the physiological role of light signaling, we next analyzed the impact of blue, red, white lights and darkness on ascorbate content in WT and *plp* mutants. One-month-old plants grown in the greenhouse were transferred to a growth chamber equipped with LEDs emitting white, blue and/or red light. Plants were first transferred to a diel cycle of 12 h white light at 250-260 µmol. m^-2^.s^-1^ and 12 h of darkness for four days. Then, at dawn, they were transferred under four light regimes, 100% blue light, 100% red light or 50% blue/40% red light, respectively, and darkness or maintained under white light as control. Light intensity was the same for all conditions, except for darkness (Suppl. Fig. 6). In the WT under white light, leaf ascorbate content was increased during the first part of the photoperiod and decreased during the second part of the photoperiod (Fig. 5A-B). In the mutant, a similar evolution was found, with a similar amplitude between minimum and maximum values, but a higher basic level and a more sustained increase during the day. The same pattern was found for the mutant under the different light regimes, but not for the WT, in which ascorbate was no longer increased under darkness and red light (Fig. 5C-F). Hierarchical clustering analysis showed a clear difference between WT and mutant, which clustered separately (Fig. 5G). Then, in the WT, two clusters were clearly distinguished, firstly darkness and red light, and secondly white, blue/red and blue light. This was not the case in the mutant, for which red, blue/red and darkness clustered together. These results suggest that blue light promotes ascorbate accumulation by counteracting the inhibitory effect of PLP on ascorbate synthesis. Given that GGP, one of the very few proteins described as interacting with PLP (Ogura et al., 2008), was found to interact *in vivo* with PLP (see above), we next investigated its effect on the activity of GGP *in vitro*.

**Figure 5.**
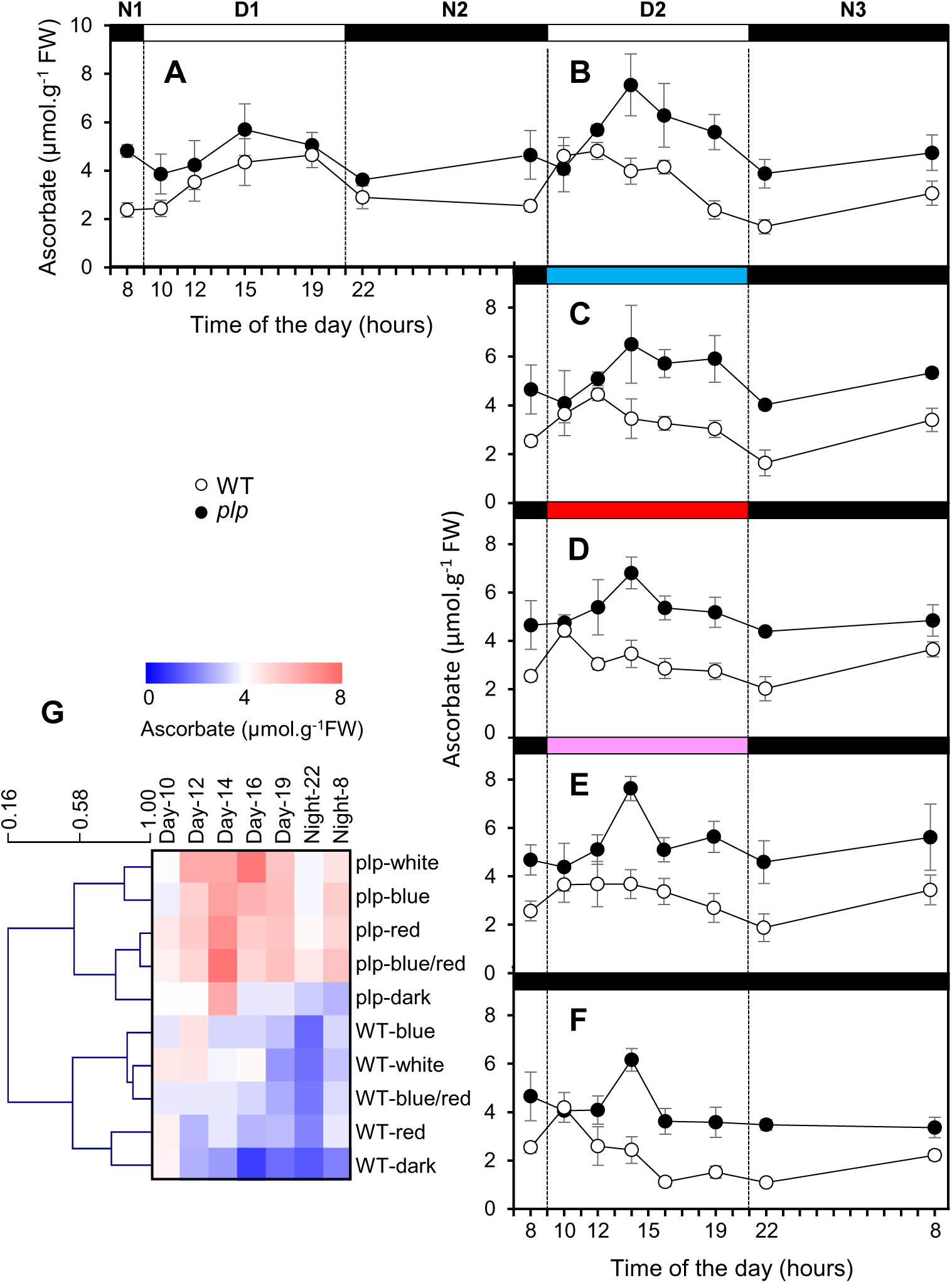
Effect of light on ascorbate evolution in WT and *plp* mutant leaves during a day-night cycle. A. In leaves of plants grown in a greenhouse then transferred to a growth chamber under a white light intensity of 260-270 µmol.m^-2^.s^-1^ for 24 hours. B. Following cycle, still under white light. C. Following cycle under blue light. D. Following cycle under red light. E. Following cycle under blue (50%) and red (40%) light. F. Following cycle in the dark. G. Heat map representing a clustering analysis performed with mean values in MEV4.9. Columns correspond to time, and lines correspond to clustered content of ascorbate based on Pearson’s correlation coefficient (XXX algorithm?). All data shown in A-F are expressed as means ± SD (n=4).

### Blue light prevents PAS/LOV inhibition of GDP-L-galactose phosphorylase activity *in vitro*

In order to study the *in vitro* interaction between PLP and GGP, the two tomato proteins were expressed heterologously. In the case of GGP, a functional protein with characteristics relatively close to those already published (Linster et al., 2008) was obtained with *E. coli*. Indeed, using GDP-alpha-glucose as a substrate, the K_m_ and specific activity were respectively 17 µM and 37 U.mg^-1^ against 4-12 µM and 10-16 U.mg^-1^ in Arabidopsis (Linster et al., 2008). In contrast, attempts to obtain a PLP capable of inhibiting GGP *in vitro* by expressing it in *E. coli* were unsuccessful, including accumulation of the protein in inclusion bodies that could only be resolubilized using chaotropic conditions. However, the latter did not make it possible to obtain a functional protein. Using a transient expression system in tobacco (Yamamoto et al., 2018) we obtained a PLP capable of inhibiting the activity of GGP (Fig. 6A). The inhibition of GGP by PLP set in quickly, probably within seconds (the time resolution of the spectrophotometer did not allow for more precise data) and was stable for hours (not shown). Heating of PLP at 95°C for 10 min suppressed this effect. Strikingly, large amounts of PLP were necessary to inhibit GGP. Thus, for a GGP concentration of 3 nM, 90% inhibition was obtained with 900 nM of PLP, *i.e.* at a ratio of about 300. A kinetic study showed that the inhibition is non-competitive (Fig. 6B), PLP reducing the maximal activity but not the affinity for the substrate (here GDP-alpha-glucose). When applied to PLP before mixing with GGP, blue light (445 ± 15 nm) counteracted its inhibitory effect at a level that depended on both the intensity and the duration of the illumination (Fig. 6C). PLP appeared sensitive to blue light, with a response already significant at 25 µmol.s^-1^.m^-2^ and a plateau reached at around 250 µmol.s^-1^.m^-2^ for the rate of inactivation, confirming that the blue light intensities used in the *in vivo* experiment described above (Fig 5) were effective. In contrast, blue light had no effect when applied after having mixed PLP and GGP (not shown). Red light (2000 µmol.s^-1^.m^-2^ at 653 ± 33 nm) also had no effect on the interaction between PLP and GGP (not shown). Finally, we tested the reversibility of the action of blue light by illuminating PLP with blue light at maximal intensity (2000 µmol.m^-2^.s^-1^) during 2 h before transfer to darkness for 6 h, the time needed to fully recover the ‘dark’ form (Ogura et al., 2007). While after 2 h, the inhibition was 86% and 16% for PLP incubated in the darkness and under blue light, respectively, it was 77% and 32% after 6 h of additional incubation in the dark. This confirms that PLP returns to its ‘dark’ active form only very slowly following its exposure to blue light (Suppl. Fig. 7) .

**Figure 6.**
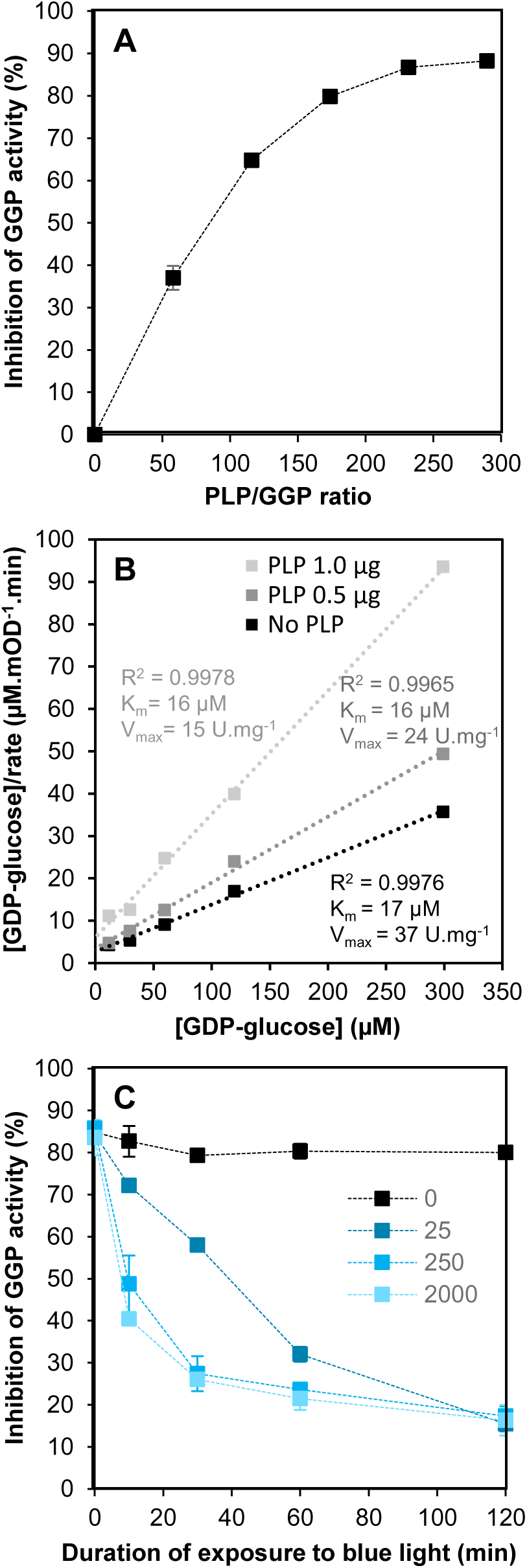
*In-vitro* inhibition of GDP-L-galactose phosphorylase by PAS/LOV and effect of blue light on PAS/LOV. A. Relation between the PLP/GGP ratio and the inhibition of GGP. B. Hanes-Woolf plot GDP-L-galactose phosphorylase inhibition by PLP. GDP-glucose at concentrations of 12, 30, 60, 120 and 300 µM was used as substrate. C. Effect of blue light exposure duration on GGP inhibition by PLP. The light was applied before mixing the two proteins. All data shown are expressed as means ± SD (n=2 technical duplicates).

## Discussion

Given the importance of ascorbate for human health and plant performance, the study of ascorbate metabolism in plants holds great interest from both agronomic and economic perspectives. Since the discovery of the main ascorbate biosynthetic pathway (Wheeler et al., 1998) and the characterization of the enzymes involved (Conklin 1999 and 2006; Linster et al., 2007), many studies have tried to decipher the regulatory mechanisms involved. Although it has long been known that light modulates ascorbate metabolism in plants, the underlying mechanisms were far from understood. In the present study, we identified a major player of the regulation of ascorbate metabolism by identifying the causal mutation leading to a 2 to 4-fold increase in the ascorbate content in an EMS Micro-Tom mutant. The recessive mutation corresponded to a stop codon in the *Solyc*05g007020 gene that encodes a photoreceptor PAS/LOV protein. Our work allowed (i) to undoubtedly define PLP as a negative regulator of ascorbate biosynthesis, (ii) to confirm the interaction of PLP with the GGP protein, (iii) to provide *in vitro* evidence that this interaction results in the inhibition of the activity of GGP and (iv) to demonstrate that blue light counteracts this inhibition.

### Why blue light as a regulator of ascorbate synthesis?

The function of PLP, which was first reported twenty years ago (Crosson et al., 2003), has remained unknown despite the demonstration of its interaction with GGP (Ogura et al., 2008) and conformational change induced by blue light (Kasahara et al., 2010). The presence of a LOV domain links it to phototropins, which have two LOV domains (LOV1 and LOV2) and initiate various responses to blue light (phototropism, opening of stomata, chloroplast movements, leaf expansion and movements) *via* autophosphorylation or even transphosphorylation (Christie, 2007). Members of the ZTL/ADO family, which, like PLP, have only one LOV domain, are involved in the photo-control of flowering and regulating the circadian clock (Krauss et al., 2009). More generally, blue light stimulates shoot compactness by repressing the growth of the hypocotyl and internodes and promoting leaf thickness, or flowering, and the production of secondary compounds such as carotenoids and flavonoids, as well as photosynthesis (Huche-Thelier et al., 2016). Plants would therefore use blue light to perceive the amount of light energy available in order to optimize their developmental program, but also to protect themselves from an excess of energy, in particular thanks to the different levels of photosensitivity of the photoreceptors (Christie, 2007). This idea is reinforced by the fact that flavins that generate ROS when excited by blue light are precisely the cofactors of these photoreceptors (Losi and Gartner, 2012). The promoting effect of blue light on ascorbate synthesis would enable it to cope with the increase in ROS production when light intensity augments. In addition, higher ROS content tends to increase the proportion of the oxidized form of ascorbate, which is less stable (Bulley and Laing, 2016; Truffaut et al., 2017). However, while phototropins are activated by blue light, leading to the transduction of this signal *via* interaction with their target proteins, the opposite is observed with PLP which, on the one hand, targets an enzyme of the pathway and, on the other hand, no longer acts or acts very weakly once exposed to blue light. PLP would therefore represent a peculiar evolution among photoreceptors, although favored by a modular nature that has allowed the appearance of numerous combinations of domains and, therefore, of neo-functionalization during evolution (Moglich et al., 2010). Strikingly, the photocycle rate is highly variable in photoreceptors containing the LOV domain. Thus, while photoexcitation only lasts a few tens of seconds in phototropins (Kasahara et al., 2002), it is maintained much longer in members of the ZTL/ADO family with more than 60 h for FKF1 (Zikihara et al., 2006), as well as in PLP, for which several hours were necessary for a return to the dark form (Kasahara et al., 2010), a result confirmed in the present work *via* the *in vitro* measurement of its inhibitory action. The irradiation of a LOV domain causes the formation, within a few microseconds, of a covalent bond between FMN and a nearby cysteine residue (Christie et al., 2015). While after 10 seconds of exposure to blue light, a significant change in the absorption spectrum of tomato PLP expressed in *E. coli* reflecting the change of state of FMN was found (Kasahara et al., 2010), several tens of minutes were necessary to reach minimum effect of PLP on GGP. However, PLP responded to low light intensity, with the inactivation rate increasing almost linearly until it plateaued at an intensity corresponding to the fraction of blue light in direct sunlight, *i.e.* about 200 µmol.m^-2^.s^-1^. These results are not necessarily contradictory if we consider that higher intensity increases the probability that PLP inactivation would occur, inactivation being different from the change of state of FMN itself. Moreover, PLP expressed in *E. coli* was not able to inhibit GGP, while that expressed in *Nicotiana benthamiana* was. It is therefore possible that a difference in folding would affect its photosensitivity.

### How blue light promotes ascorbate synthesis in a diel cycle in leaves

The results obtained here introduce an additional layer of complexity to the regulation of ascorbate metabolism in general, and GGP in particular. The latter, admitted as being the most controlling enzyme of the ascorbate synthesis pathway (Fenech et al., 2021), already known to be regulated by a range of factors, including light and stress, at the transcriptional (Bulley and Laing, 2016) and translational (Laing et al., 2015) levels, appeared to be also regulated post-translationally. This also extends the list of processes associating ascorbate synthesis and light, such as those involving the AMR1 protein (Zhang et al., 2009), and the COP9 signalosome complex (Mach 2013). It is striking that the dark form of PLP formed a stable complex with GGP that could hardly be dissociated by blue light (Fig. 7). The latter would therefore only act on newly formed copies of PLP, preventing them from interacting with GGP. The timing and amplitude of the expression of these two proteins would therefore be essential to condition their interaction. Figures 3 and 5 show that ascorbate exhibited relatively large fluctuations during day-night cycles and that the loss of PLP amplified them. In both cases, ascorbate increased during the day, dropped at the end of the day or the beginning of the night, increased again during the night, then dropped again at the beginning of the day. These fluctuations can be attributed to the amount of substrate available for ascorbate synthesis, the rate of ascorbate turnover, and the amount of active GGP (Bulley and Laing, 2016). The expression of genes encoding GGP1 (in tomato by far the most expressed of the two isoforms of this enzyme) and PLP increased at night to both peaks towards the end of the night. Interestingly, GGP1 expression was lower in the mutant (Fig. 5), a result that seems to confirm that ascorbate exerts a negative feedback on GGP expression but contradicts the absence of effect of ascorbate itself reported in Arabidopsis (Laing et al., 2015; Bulley et al., 2021), suggesting that another factor could be at play. The fact that ascorbate increased indicates that there were not enough copies of PLP to neutralize all those of GGP formed overnight. Thereafter, GGP expression dropped to zero or very low for most of the day, while PLP expression was retained and even increased during the day. Consequently, the number of copies of PLP would then become sufficient to block GGP, as evidenced by the decrease in ascorbate observed in the WT, but not in the mutant during the photoperiod under red light. In contrast, this is not true under white or blue light under which PLP would be deactivated (Fig. 5). Taken together, these elements suggest that the stoichiometry between GGP and PLP plays an essential role, the interaction between PLP and blue light making it possible to adjust the production of ascorbate to light intensity and ultimately to the production of light-dependent ROS. Therefore, it will be interesting to study the turnover of both proteins in diel cycles and under various light regimes, but also in other organs, in particular fruit, in which PLP also modulates ascorbate synthesis (Fig. 2). In the meantime, it seemed interesting to investigate whether transcriptomic and proteomic data would be available for these two genes.

**Figure 7.**
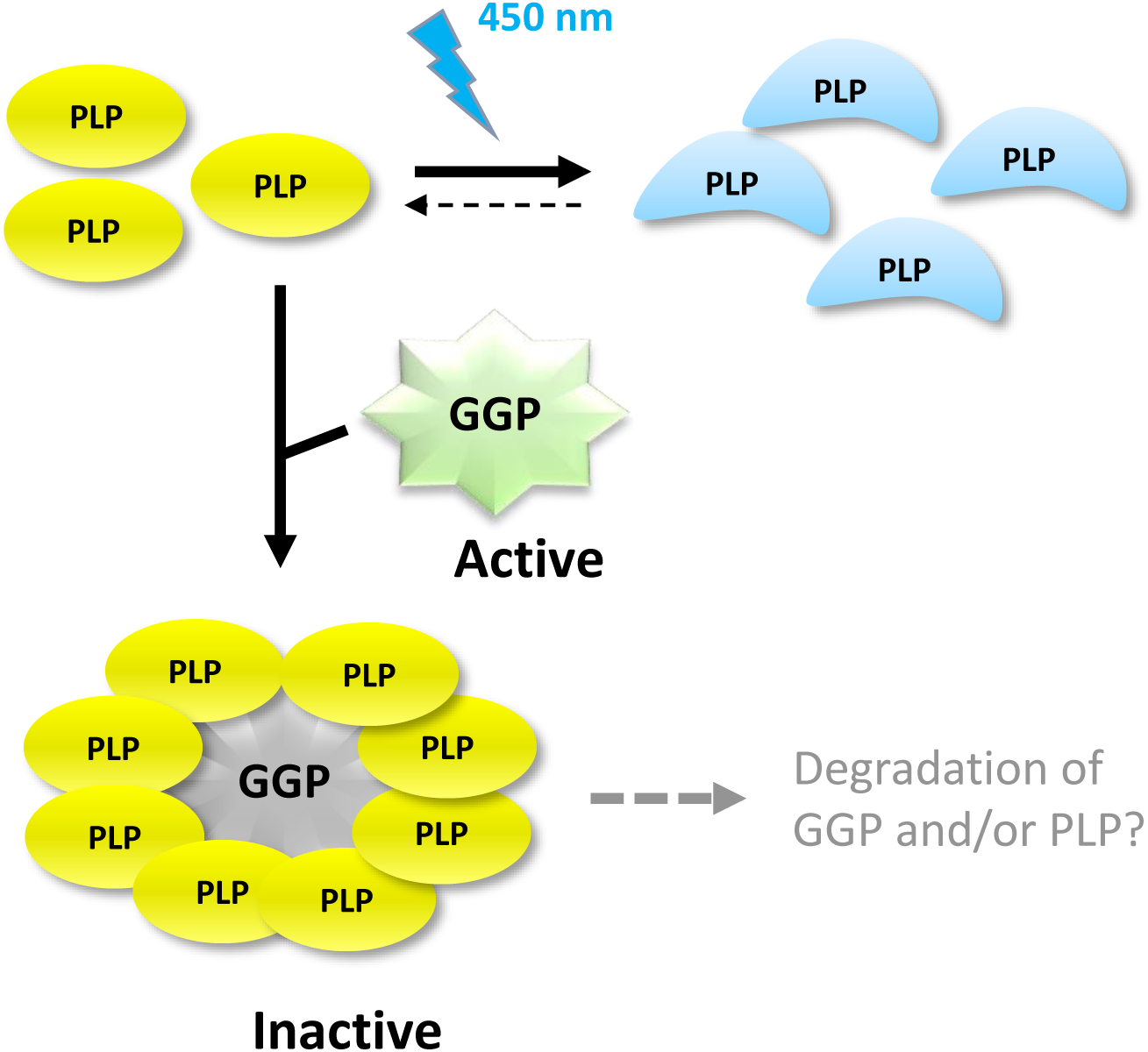
Schematic model describing the activation of ascorbate synthesis by blue light. Newly synthesized PAS/LOV protein binds GDP-L-galactose phosphorylase unless deactivated by blue light. Its deactivated form is stable for several hours while its active form irreversibly inhibits its target, possibly leading to its degradation. Abbreviations: PLP, PAS/LOV; GGP, GDP-L-galactose phosphorylase.

### Regulation of ascorbate synthesis by PAS/LOV as an unusual and costly mechanism

Available transcriptomic data indicate that the expression levels of GGP and PLP are elevated (Suppl. Fig. 8A-B). Thus, in the Arabidopsis leaf during a day-night cycle (Bläsing et al., 2005), VTC2 was among the 2-3% and PLP among the 10-30% of the most highly expressed genes based on their average expression. Interestingly, the expression of PLP was strongly stimulated during the night in a starchless mutant, probably related to carbon starvation occurring at night in the mutant (Bläsing et al., 2005). Indeed, this gene is one of the genes most strongly repressed by sucrose and glucose (Bläsing et al., 2005). Yet, neither PLP nor VTC2 (GGP1) was found among the almost 5000 proteins recently detected in a day-night cycle in the leaves of the same Arabidopsis accession (Uhrig et al., 2021). Similar results were found during tomato fruit development (Belouah et al., 2020), where the expression levels of GGP1 and PLP were also high, being respectively among the 2% and 22% of the most highly expressed genes based on the average calculated for the entire development (Suppl. Fig. 8C-D). The encoded proteins were still not found among more than 2800 proteins detected. These proteins were also not present in a larger dataset of almost 8000 proteins detected in tomato fruit (Szymanski et al., 2017). These observations are supported by the fact that the activity of GGP measured in the leaves of Arabidopsis is very low (Yoshimura et al., 2014). Among the explanations for such a discrepancy, one could invoke ’inefficient’ translation, in particular for GGP whose translation is jointly repressed by uORF and ascorbate (Laing et al., 2015), and/or high instability of the protein. It therefore seems relevant to ask whether the interaction between the two proteins leads to their degradation. Indeed, we found *in vitro* that the interaction was stable for at least several hours and could not be reversed even by the application of very intense blue light. The ratio of 300 found between PLP and GGP to block the activity of the latter may be overestimated, given that the proportion of functional protein among purified PLP was not known. In order to learn more about the stoichiometry between these two proteins, it will be interesting to carry out crystallography studies. Apart from blocking copies of PLP but also of GGP, this suggests that the interaction *in vivo* between GGP and PLP could lead to the degradation of GGP or even that of PLP. The dual localization between the cytosol and the nucleus observed by microscopy could be linked to the turnover of these proteins. Indeed, LKP2, a member of the ZTL/ADO family located in the nucleus, has been shown to form a complex that functions as a ubiquitin E3 ligase and interacts with the circadian clock (Yasuhara et al., 2004), probably by addressing one or more components of the latter to the 26S proteasome. It would therefore be interesting to search for a similar mechanism that would involve PLP and GGP. It is also worth mentioning that two members of the TALE homeodomain protein transcription factor family, which is involved in numerous developmental processes (Hackbusch et al., 2005), have been identified as interacting with PLP by a double-hybrid screening of an Arabidopsis cDNA library (Ogura et al., 2008).

Finally, the mechanism regulating ascorbate synthesis shown here appears to be very costly with respect to GGP expression. However, the very high expression of GGP can also be seen as providing a great flexibility in response to stress. For example, an increase in sugars, often observed in response to various stresses, would repress the expression of PLP, thus making it possible to quickly increase the activity of the most controlling enzyme of the ascorbate synthesis pathway. These valuable data on the regulation of ascorbate metabolism will also make it possible to envisage new strategies for improving the nutritional quality of cultivated plants and their ability to withstand environmental stresses.

## Materials and Methods

### Plant material and culture conditions

*Solanum lycopersicum* L. cv. Micro-Tom was used for all experiments performed in this study. Plant culture conditions in greenhouse were as in Rothan et al. (2016). To study the effect of light on ascorbate, experiments were carried out in spring or summer with one-month old plants grown in the greenhouse. Thereafter, the plants remained either in the greenhouse in the case of the analysis of the ascorbate change in the time course of fruit development or the whole plant organs. For the “light assay”, the harvesting time of the plant materials was designed according to the light period (https://jekophoto.eu/tools/twilight-calculator-blue-hour-golden-hour/). This experiment was carried out in perfect weather conditions two times, on May 19th and July 10th, 2018. One-month-old plants cultured in the green house were moved out the night before the beginning of the experiment, the starting time was at 10 am, and every two hours 3 leaves per plant on three plants were harvested. At the same time temperature, hygrometry, photosynthetically active radiation (PAR, in µmol of photons m^-2^s^-1^) were measured with a Quantum photometer with a light sensor LI-190R (LI-COR Corporate, Lincoln Nebraska, USA) while the light spectrum was recorded with a JAZ Spectrometer (Ocean Optics Inc, Largo, Florida, USA).

To investigate the effect of the light spectrum, one-month-old plants were cultured in growth chambers (HiPoint Bionef Montreuil, France) as illustrated in Suppl. Figure 6. Moving plants from the greenhouse to the growth chambers was done at night to avoid any stress. Then, the plants were acclimatized for three days under a 12 h photoperiod of white light with a photosynthetic photon flux density of around 260-265 µmol photons m^-2^ s^-1^ at 10 cm of the LED source and corresponding to the height of top leaves. The day/night temperature and relative hygrometry were 25/20°C and 65-60%, respectively. The first hour of the photoperiod (9 am to 10 am) was programmed as a ramp resulting to a linear increase of the light intensity up to the set-up, and reversely at night to a decrease between (8 pm to 9 pm) to the return of darkness, in a way to mimic sunrise and sunset, respectively. The fourth day, the same culture conditions were maintained, and the first harvesting time of 3 leaves per plant on four plants was at 8 am, just before the beginning of the photoperiod and then every two to three hours. On the fifth day, the plants were submitted to four light conditions, white (as control), blue, red, blue and red, and darkness, and the leaves were harvested up to the end of the next night. For each light condition, the photon flux density was maintained at 260-270 µmol photons m^-2^ s^-1^ (Suppl. Fig.6).

For the transient transformation experiments by agroinfiltration, WT *Nicotiana benthamiana* L. plants were cultured in soil during one month in the greenhouse before being used. Watering was carried out three times per week, but only once with a Liquoplant fertilizing solution (1.85 g L-1, Plantin SARL Courthezon, France) and twice using tap water at pH 6. For the BiFC (Bimolecular Fluorescence complementation) analyses by confocal microscopy, freshly harvested onion *Allium cepa* was purchased from the local market.

To determine ascorbate content, leaves or fruits at several stages of development were collected, cut into small pieces and immediately quenched into liquid nitrogen. Samples were stored at -80°C until extraction.

### Mapping-by-sequencing

Mapping-by-sequencing was performed as described in Garcia et al. (2016). A mapping BC_1_F_2_ population of 440 plants was created by crossing the P21H6 cv Micro-Tom ascorbate-enriched mutant line with a WT cv Micro-Tom parental line. Two bulks were then constituted by pooling 44 plants displaying either a mutant fruit phenotype (AsA+ bulk) or 44 plants with a WT phenotype (WT-like bulk). To this end, 25 leaf discs (5 mm diameter each) were collected from each BC_1_F_2_ plant (approximately 60 mg fresh weight) and pooled into the AsA+ bulk and in the WT-like bulk. The same amount of plant material was also collected from the WT parental line. Genomic DNA was extracted from each bulk and the parental line using a cetyl-trimethyl-ammonium bromide method as described by Garcia et al. (2016). DNA was suspended in 200 mL of distilled water and quantified by fluorometric measurement with a Quant-it dsDNA assay kit (Invitrogen). Illumina paired-end shotgun-indexed libraries were prepared using the TruSeq DNA PCRF-Free LT Sample Preparation Kit according to the manufacturer’s instructions (Illumina). The libraries were validated using an Agilent High Sensitivity DNA chip (Agilent Technologies) and sequenced using an Illumina HiSeq 2000 at the INRA EPGV facility, operating in a 100-bp paired-end run mode. Raw fastq files were mapped to the tomato reference genome sequence *S. lycopersicum* build release SL3.0 (ftp://ftp.solgenomics.net) using BWA version 0.7.12 (Li and Durbin, 2009; http://bio-bwa.sourceforge.net/). Variant calling (SNPs and INDELs) was performed using SAMtools version 1.2 (Li et al., 2009; http://htslib.org). As the tomato reference genome (cv Heinz 1706) used to map the reads is distinct from that of cv Micro-Tom, the variants identified would include both cv Heinz 1706/cv Micro-Tom natural polymorphisms in addition to EMS mutations. In this context, additional sequencing to a minimum depth of 203 of the cv Micro-Tom line was performed to take into account and further remove the cv Heinz 1706/cv Micro-Tom natural polymorphism. The output file included various quality parameters relevant to sequencing and mapping that were subsequently used to filter the variants. The cv Micro-Tom line output file (.vcf) included all variants (SNPs plus INDELs) corresponding to natural polymorphisms between cv Micro-Tom and cv Heinz 1706. The two .vcf output files obtained from the AsA+ and WT-like bulks included variants (SNPs plus INDELs) corresponding to natural polymorphisms between cv Micro-Tom and cv Heinz 1706 and also to EMS mutations. The .vcf files were annotated using SnpEff version 4.1 (http://snpeff.sourceforge.net/SnpEff.html; Cingolani et al., 2012) using ITAG2.40 gene models (ftp://ftp.solgenomics.net) SNP allelic frequencies between AsA+ and WT-like bulks and the cv Micro-Tom parental line were compared using a custom Python script version 2.6.5 (https://www.python.org). Once the putative causal mutation was detected using the mapping-by-sequencing procedure, the EMS-induced SNPs flanking the putative mutation were used as markers for genotyping the BC_1_F_2_ individuals using a KASP assay (Smith and Maughan, 2015). Specific primer design was performed using batchprimer3 software (Smith and Maughan, 2015; http://probes.pw.usda. gov/batchprimer), and genotyping was done using KASP procedures (LGC Genomics).

### CRISPR/Cas 9 gene editing of PLP and stable tomato transformation

CRISPR/Cas9 gene editing was performed as described in Fauser et al. (2014). A construction comprising a single sgRNA alongside the Cas9 endonuclease gene, was designed to induce target deletions in PAS/LOV-coding sequence. The sgRNA target sequence was designed using CRISPR-P 2.0 web software (http://crispr.hzau.edu.cn/CRISPR2/; Lei et al., 2014). Since targeting-RNAs are inserted into pDECAS9 vector, the final plasmid was used to transform Micro-Tom tomato cotyledons through Agrobacterium infection as described in Fernandez et al. (2009). The T0 plants resulting from the regeneration of the cotyledons were genotyped and their fruits were phenotyped for ascorbate content. T1 seeds from selected ascorbate-enriched T0 plants were sown for further characterization. The CRISPR/Cas9 positive lines were further genotyped for indel mutations using primers flanking the target sequence. To obtain tomato plants overexpressing the GFP-PLP fusion protein, the *Pro35S:eGFP-PLP* construct in pK7WGF2 vector was used for the transformation as described above. All primers used for cloning are shown in Suppl. Table 5.

### Ascorbate assay

Samples were ground to a fine powder using a TissueLyser II (Qiagen). Ascorbate content was assayed using a protocol adapted from Bergmeyer (1987) using 40 and 100 mg FW for leaves and fruits, respectively, extracted in 400 µL 0.1 M HCl. For the determination of total ascorbate, 20 µL of extract were first incubated 10 min in 0.15 M HEPES/KOH pH 7.5 and 0.75 mM DTT. N-Ethyl maleimide was added to reach 0.035% w/v. After 10 min, 1 unit.mL^-1^ of ascorbate oxidase was added. After 20 min, phenazine ethosulfate and thiazolyl blue mix were added for final concentrations of respectively 0.3 mM and 0.6 mM. The thiazolyl blue mix was prepared as following: 10 mM thiazolyl blue, 0.2 M Na_2_HPO_4_, 0.2 M citric acid, 2 mM EDTA and 0.3% v/v Triton X100 at pH 3.5. All steps were performed in a polystyrene microplate and at room temperature. Measurements were performed at 570 nm in MP96 readers (SAFAS, Monaco). For reduced ascorbate, the same protocol was used, except steps involving DTT and N-ethylmaleimide were omitted.

### RT–qPCR analysis

Total RNA was extracted from leaves using Trizol reagent (Invitrogen) and purified with a RNeasy Plant Mini Kit (Qiagen). Relative transcript levels were determined as described previously (Deslous et al. 2021) using gene-specific primers and *eIF4A* and *β-tubulin* as an internal control. The primer sequences are shown in Suppl. Table 5.

### Protein subcellular localization

All constructs used were realized using gateway® technology (Invitrogen). The cDNA without STOP codon (NS) of GGP1, GGP2, PLP and PLP^m^ were synthesized by GeneArt® Gene Synthesis (Invitrogen), and directly provided into entry vector pDONR201™ (Suppl. Table 5). The mutation of PLP construct was the same as the one identified in EMS mutant. In order to obtain fusion proteins with fluorescent tag either in C-terminal or N-terminal, specific primers (listed in Suppl. Table 6) was used to add a STOP codon and the flanking AttB sequences by PCR reaction. Classical BP recombination reactions allowed to insert the new sequences into a pDONR201™ and then LR reaction permitted to transfer our sequence of interest into the different destination vectors. The *Agrobacterium tumefaciens* electro-competent strain GV3101 was transformed with the above fluorescent fusion constructs. Transformed agrobacteria were selected on LB medium supplemented with suitable antibiotics and conserved at -80°C. Prior to agroinfiltration, inoculated LB cultures were incubated overnight at 28°C until 0.6 to 0.8 OD_600nm_. For subsequent infiltration, the culture was centrifuged and the pellet suspended in water to reach 0.2 OD_600nm_ in the case of sub-cellular localization. Then, 50-100µL of this bacterial solution was infiltrated in the leaf epidermis of three-week-old *N. benthamiana* plants at the level of a wounding by needle using a 1 mL syringe to improve infiltration. The plants were maintained in normal culture conditions (light, temperature) for 48h and the observation was carried out on the underside of the leaf epidermis.

### Protein interactions in plant cell

To assess the interaction of PLP and GGP1 in plant cell, a BiFC (Bimolecular Fluorescence Complementation) (Walter, 2004) experiment was performed in onion epidermal cells by biolistic transformation. Each cDNA of the genes of interest were inserted using the GATEWAY technique in different vectors in order to test all possible orientations of the protein fusions (Suppl. Table 5). Then, 2.5 μg of plasmid DNA of each construct were mixed with 25 μl of a suspension (250 μg / μl) of gold micro-particles (diameter = 0.6 μm) in 50% ethanol (v / v) then 25 μl of 2.5 M CaCl_2_ and 10 μl of 0.1 M spermidine are added. The micro-particles were let to sediment for 10 min before being washed with 70% and 100% (v/v) ethanol. Eight microliters of the micro-particle suspension (30 μl) were used for transformation of epidermal onion cells using the PDE-1000He particle gun (Bio-rad). Before transformation, the onion epidermis was taken from the innermost scales of the bulb and deposited, upper face in contact with MS medium. Transformation of onion epidermis cells was performed at a pressure of 710 mm Hg at a helium pressure of 1100 psi and at a distance of 6 cm.

### Imaging

Live imaging was performed in the plant imaging division of the BIC platform (Bordeaux Imaging Centre), using a Zeiss LSM 880 confocal laser scanning microscopy system equipped with 40x objectives. The excitation wavelengths used for the eGFP (or YFP) and mCherry were 514 and 543 nm, respectively. The emission windows defined for their observation were respectively between 525 and 600 nm for the eGFP (or YFP) and between 580 and 650 nm for the mCherry.

### Optogenetic assay

In this animal cell system, human embryonic kidney cells, namely HEK-293T, were transfected by a combination of plasmids (Suppl. Table 5) and cultured as described by Müller et al. (2013). For the experimental set-up, 50,000 cells were seeded into 24-well plates. 24 h after seeding cells were transfected using a polyethylenimine-based (PEI, linear, MW: 25 kDa, Polyscience) method, as described elsewhere (Müller et al., 2013). If co-transfected, plasmids were applied in equal -weight-based-amounts. Four hours post transfection, the cell-culture medium was exchanged by prewarmed fresh medium under green safelight conditions. 20 h later, cells were illuminated, using LED-panels emitting blue light of a wavelength of 455 nm for 24 h (reporter gene activity) while control cells were kept in the dark. The quantitative determination of the activity of the secreted alkaline phosphatase (SEAP) in the cell culture medium was performed by using a previously described colorimetric assay (Müller et al., 2014; Beyer et al., 2015).

### Proteomic analysis

Protein samples from tomato leaves were prepared by grinding 200 mg of leaves in 1 mL of 2x Laemmli buffer for 5 min in liquid nitrogen. Proteins were further solubilized by heating at 80°C for 20 min. The insoluble material was removed by centrifugation (20 min at 13,000g), and the proteins of the supernatant were separated by SDS-PAGE (Laemmli, 1970) using 4% stacking gel and 10% running gel. After colloidal blue staining, 3 bands were cut out from the SDS-PAGE 10% gel and subsequently cut in 1 mm x 1 mm gel pieces. Gel pieces were destained in 25 mM ammonium bicarbonate 50% acetonitrile, rinsed twice in ultrapure water and shrunk in acetonitrile for 10 min. After acetonitrile removal, gel pieces were dried at room temperature, covered with the trypsin solution (10 ng/µL in 40 mM NH_4_HCO_3_ and 10% acetonitrile), rehydrated at 4 °C for 10 min, and finally incubated overnight at 37°C. Spots were then incubated for 15 min in 40 mM NH_4_HCO_3_ and 10% acetonitrile at room temperature with rotary shaking. The supernatant was collected, and an H_2_O/acetonitrile/HCOOH (47.5:47.5:5) extraction solution was added onto gel slices for 15 min. The extraction step was repeated twice. Supernatants were pooled and concentrated in a vacuum centrifuge to a final volume of 30 µL of 0.01% HCOOH. Digests were finally stored at -20 °C.

The peptide mixture was analyzed on an Ultimate 3000 nanoLC system (Dionex, Amsterdam, The Netherlands) coupled to an Electrospray Q-Exactive quadrupole Orbitrap benchtop mass spectrometer (Thermo Fisher Scientific, San Jose, CA). Ten microliters of peptide digests were loaded onto a 300-µm-inner diameter x 5-mm C18 PepMapTM trap column (LC Packings) at a flow rate of 30 µL/min. The peptides were eluted from the trap column onto an analytical 75-mm id x 25-cm C18 Pep-Map column (LC Packings) with a 4–40% linear gradient of solvent B in 108 min (solvent A was 0.1% formic acid in 5% acetonitrile, and solvent B was 0.1% formic acid in 80% acetonitrile). The separation flow rate was set at 300 nL/min. The mass spectrometer operated in positive ion mode at a 1.8-kV needle voltage. Data were acquired using Xcalibur 2.2 software in a data-dependent mode. MS scans (m/z 350-1600) were recorded at a resolution of R = 70 000 (@ m/z 200) and an AGC target of 3 x 106 ions collected within 100 ms. Dynamic exclusion was et to 30 s and top 12 ions were selected from fragmentation in HCD mode. MS/MS scans with a target value of 1 x 105 ions were collected with a maximum fill time of 100 ms and a resolution of R = 17500. Additionally, only +2 and +3 charged ions were selected for fragmentation. Others settings were as follows: no sheath nor auxiliary gas flow, heated capillary temperature, 250 °C; normalized HCD collision energy of 25% and an isolation width of 2 m/z.

### Heterologous expression and purification of PLP and GGP1

The codon-optimized PLP was chemically synthesized by Proteogenix (Schiltigheim, France) and then cloned into the pET32a vector (Novagen). The PLP was expressed as a fusion protein with thioredoxin and His_6_ at the N-terminus in *E. coli* BL21(DE3)pLysS (Novagen) host cell. A bacterial culture was performed as described in Kasahara et al. (2010). The BL21(DE3)pLysS transformant harboring the PLP construct was grown at 25°C in M9 medium supplemented with ampicillin (100 µg mL^-1^) and chloramphenicol (25 µg mL^-1^). When the culture reached 0.35 OD_600nm_, the PLP expression was induced in the presence of 1 mM isopropyl β-D-thiogalactopyranoside for 18 h at 25 °C. The cells were harvested by centrifugation, the pellet suspended in 30 mL of lysis buffer (Tris-HCl 50 mM pH8, NaCl 0.5 M, glycerol 2% (v/v), 5 mM β-mercaptoethanol, 0.2% Sarkosyl, imidazole 50 mM, protease inhibitor cocktail EDTA free (Roche), lysozyme 100 µg mL^-1^) keep at room temperature for 20 min. Then, the suspension was frozen in nitrogen liquid and thaw at 25°C twice, before sonication for 15 min on ice. The lysate was centrifuged at 30,000g for 30 min at 4 °C, and the supernatant was filtered on 0.45 µm units before loading onto a nickel-Sepharose Fast Flow column (1mL of bed volume) using an ÄKTA^TM^ Start Fast Protein Liquid Chromatography system (GE Healthcare). The PLP protein was eluted at 160 mM imidazole. The fractions containing the PLP peak were pooled and desalted on PD10 column equilibrated with HEPES/KOH 50mM pH7.5, glycerol 2 % and Sarkosyl 0.2 % (w/v) 5 mM β−mercaptoethanol, then concentrated using Vivaspin® 6 concentrators (Sartorius Stedim Lab Ltd, UK) and stored at -80 °C before use.

The full-length coding sequence of GGP1 was amplified by PCR and cloned into the pET28a vector (Novagen). The GGP1 enzyme was expressed in *E. coli* BL21(DE3)pLysS strain as a fusion protein with His_6_ at the N terminus. The BL21(DE3)pLysS transformant harboring the GGP1 construct was grown at 37°C in LB (500 mL) medium supplemented with kanamycin (50 µg mL^-1^) and chloramphenicol (25 µg mL^-1^). When the culture reached 0.5 OD_600nm_, the GGP1 expression was induced in the presence of 1 mM isopropyl β-D-thiogalactopyranoside for 6 h at 37 °C. The protein extraction and purification were performed as described above without adding β−mercaptoethanol and Sarkosyl in the lysis and desalting elution buffers.

### PLP expression in Tobacco and purification

The expression of PLP in Tobacco plants was carried out as described by Yamamoto et al. (2018). Briefly, the full-length coding sequence of PLP with His_6_ at the N terminus was amplified by PCR and inserted into SalI-digested pBYR2HS vector using the In-Fusion Snap Assembly Master Mix (Takara Bio). The leaves of four-week-old *N. benthamiana* plants were infiltrated with the *Agrobacterium tumefaciens* GV3101, harboring pBYR2HS-PLP with OD_600nm_ adjusted approximately at 0.5. Once infiltrated, the plants were sprayed with a 200 mM ascorbate solution containing 0.1% Triton X-100 as described by Nosaki et al. (2021). At last, the plants were grown under 16 h photoperiod of red light with a photosynthetic photon flux density of around 260-265 µmol photons m^-2^ s^-1^ in a growth chamber with a day/night temperature and relative hygrometry of 25/20 °C and 65-60 %; respectively. After four days, whole infiltrated leaves were harvested and stored at -80 °C until protein extraction.

The infiltrated *N. benthamiana* leaves were ground using mortar and pestle in liquid nitrogen. All the following steps were carried out at 4 °C. Five g of leaf powder were homogenized in 40 mL of extraction buffer (50 mM Na-phosphate pH 7.5, 0.5 mM NaCl, 2% glycerol (v/v), 0.2% Tween 20 (v/v), 50 mM imidazole and 5 mM β-mercaptoethanol) using a Polytron PT2100 for 20 s. The homogenate was centrifuged at 30,000g for 30 min, and the supernatant was clarified using 0.45 µm filters before loading onto a nickel-Sepharose Fast Flow column (5 mL of bed volume) using an ÄKTA^TM^ Start Fast Protein Liquid Chromatography system (GE Healthcare). The column was washed with 40 mL of extraction buffer, followed by 15 mL of extraction buffer with 90 mM imidazole. Proteins were then eluted with a linear imidazole gradient (0.09–0.5 M). The 3 mL-fractions were analyzed by 10% SDS-PAGE gel electrophoresis, those containing the protein of interest were desalted on PD10 column equilibrated with 50 mM HEPES/KOH pH 7.5, 2% glycerol (v/v) and 0.2% Tween 20 (v/v), then concentrated using Vivaspin® 6 concentrators (Sartorius Stedim Lab Ltd, UK) and stored at -80°C before use.

### Assay of GDP-L-galactose phosphorylase activity

For the determination of GGP activity under substrate-saturating conditions, a continuous assay was used in which GGP extract was incubated at 25°C in the presence of 50 mM HEPES/KOH pH 7.5, 10 mM MgCl_2_, 2 mM EDTA, 1 mM GDP-glucose, 20 mM phosphate, 2 mM phosphoenolpyruvate, 0.5 mM NADH, 1 unit.mL^-1^ pyruvate kinase and 1 unit.mL^-1^ lactate dehydrogenase. Changes in absorbance were measured at 340 nm in a filter-based MP96 microplate reader (SAFAS, Monaco) until rates were stabilized.

The effect of light was tested by incubating purified PLP in 50 mM HEPES/KOH pH 7.5, 0.2% (v/v) Tween 20, 2% (v/v) glycerol and 1 µM FMN, under a LedHUB light source (Omicron Laserage, Laserprodukte GmbH, Rodgau-Dudenhofen, Germany) emitting at 445 ± 15 nm or 653 ± 33 nm, at intensities ranging from 0 to 2000 µmol.m^-2^.s^-1^.

### Database search and results processing

Data were searched by SEQUEST through Proteome Discoverer 2.2 (Thermo Fisher Scientific Inc.) against the *Solanum lycopersicum* protein database downloaded from the SGN website (version ITAG3.2; 35768 entries) in which the sequences of the 3 constructs were added. Spectra from peptides higher than 5000 Da or lower than 350 Da were rejected. The search parameters were as follows: mass accuracy of the monoisotopic peptide precursor and peptide fragments was set to 10 ppm and 0.02 Da respectively. Only b- and y-ions were considered for mass calculation. Oxidation of methionines (+16 Da) was considered as variable modification and carbamidomethylation of cysteines (+57 Da) as fixed modification. Two missed trypsin cleavages were allowed. Peptide validation was performed using Percolator algorithm (Käll et al., 2007) and only “high confidence” peptides were retained corresponding to a 1% False Positive Rate at peptide level.

## Supplemental Material

Supplemental Figure 1. Screening of the EMS Micro-Tom population.

Supplemental Figure 2. Identification of the *plp* mutation responsible for the ascorbate-enriched fruit phenotype by Mapping-by-sequencing.

Supplemental Table 1. Illumina sequencing of BC_1_F_2_ bulk individuals displaying an ascorbate-enriched mutant fruit or a WT-like fruit.

Supplemental Table 2. Number of SNPs in the mutant and the WT-like bulks for the P21H6-3 mutant.

Supplemental Figure 3. CRISPR Cas9 strategy for the PLP gene.

Supplemental Table 3. Ascorbate content in rep ripe fruits of T0 and T1 CRISPR *plp* plants.

Supplemental Figure 4. Subcellular localization of PLP in tomato and proteomic analysis of the leaf extract.

Supplemental Figure 5. *in vivo* protein-protein interaction of PLP and GGP1.

Supplemental Table 4. Result of the different combinations tested for the BiFC experiments between GGP1 and PLP or PLP^m^.

Supplemental Table 5. Set of oligos and plasmids used in this study.

Supplemental Figure 6. Pictures of tomato plants cultured in growth chamber under different light regimes and the corresponding light spectra measured at INRAE Bordeaux.

Supplemental Figure 7. Reversibility of the effect of blue light on GDP-L-galactose phosphorylase (GGP) inhibition by PAS/LOV (PLP).

Supplemental Figure 8.Evolution over time of transcripts encoding GDP-L-galactose phosphorylase (GGP) and the PAS/LOV protein.

## Acknowledgments

The authors are grateful to C. Cheniclet and L. Brocard from the Bordeaux Imaging Center for their assistance in the cytological analysis, S. Claverol from the Plateforme Protéome of the Centre Génomique Fonctionnelle Bordeaux for the proteomic analysis, and all the master students, C. Cerruti, S. Barré, M. Alonso, H. El Ouarrat, R. Gomez, D. Taillis, J. Hunziker, J. Paillassa and M. Zion who helped in the screening and the phenotypic characterization of the mutants. A special thanks to I. Atienza who took care of the plants during all these years. L. Lejay is also acknowledged for the scientific discussion in the framework of the INRAE BAP RARE project.

## Author Contributions

C.Bo. screened the EMS population and contributed to the identification of the causal mutation. P.D. validated the candidate gene and performed microscopy experiments. T.B. performed the optogenetic analysis. K.Mo., J.-P.M. and J.J. performed the generation of CRISPR lines and the stable transgenic line. S.G. performed the transcriptional analysis. C.Br. and L.F. performed the genetic analysis. S.C. and L.Be. performed the LC-MS proteomic analysis. K.Mo., K. Mi., L.Ba. and P.B. performed the protein expression and purification experiments. C.C., M.M. and Y.G. performed the enzymatic analysis. G.D., C.F. and D.J. participated in the set-up of the plant culture and growth chambers. C.Bo., K.Mo., P.D., Y.G. and P.B. wrote the manuscript with input from the other authors. P.P. contributed to the scientific input and English editing of the manuscript. All authors read and approved the manuscript.

